## Supplementary material for "Recurrent connectivity supports higher-level visual and semantic object representations in the brain"

#### Supplementary Figures:

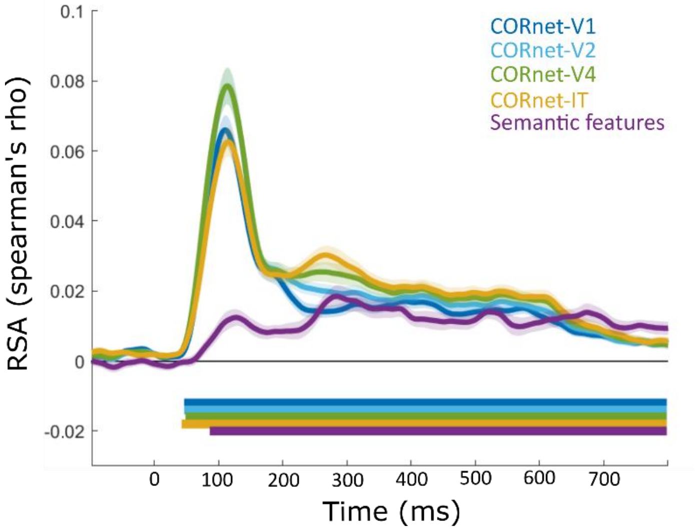


Figure S1. RSA results for the MEG sensor array using non-partial correlation RSA analysis for each model RDM over time. Shaded areas show standard error of the mean. Solid bars show time periods of significant effects.

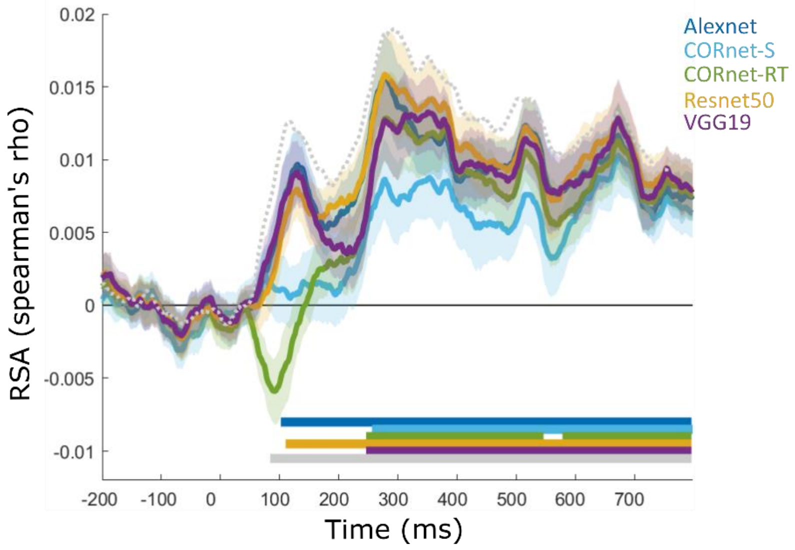


Figure S2. RSA results for the semantic feature model after different ANNs have been partialled out. Shaded areas show standard error of the mean and dotted line shows the non-partial RSA results for the semantic feature model.

### Supplementary tables:

Table S1. fMRI searchlight results

| *Model RDM* | *HO region* | *p(FWE)* | *k* | *T* | *voxel p* | *mm mm mm* |
|---|---|---|---|---|---|---|
| CORnet-V1 | left inferior lateral occipital | 0.0000 | 4497 | 11.981 | 0.0000 | [-48;-70;-1] |
| | left temporal occipital fusiform | | | 11.042 | 0.0000 | [-39;-49;-20] |
| | right inferior lateral occipital | | | 9.909 | 0.0000 | [33;-85;-1] |
| CORnet-V2 | left inferior lateral occipital | 0.0000 | 6118 | 12.553 | 0.0000 | [-45;-67;-5] |
| | right inferior lateral occipital | | | 11.940 | 0.0000 | [33;-85;-1] |
| | left temporal occipital fusiform | | | 10.491 | 0.0000 | [-39;-49;-20] |
| | right precentral | 0.0018 | 177 | 5.834 | 0.0000 | [36;-7;44] |
| | right postcentral | | | 5.208 | 0.0001 | [33;-34;51] |
| | right superior parietal | | | 5.200 | 0.0001 | [36;-55;51] |
| CORnet-V4 | left inferior lateral occipital | 0.0000 | 7352 | 12.148 | 0.0000 | [-48;-73;-1] |
| | right inferior lateral occipital | | | 11.796 | 0.0000 | [36;-82;3] |
| | right temporal occipital fusiform | | | 9.697 | 0.0000 | [33;-49;-9] |
| | left postcentral | 0.0137 | 121 | 6.161 | 0.0000 | [-54;-25;44] |
| | left supramarginal | | | 5.188 | 0.0001 | [-51;-34;33] |
| CORnet-IT | left inferior lateral occipital | 0.0000 | 5711 | 10.529 | 0.0000 | [-45;-70;-1] |
| | right inferior lateral occipital | | | 8.870 | 0.0000 | [45;-76;-13] |
| | right inferior lateral occipital | | | 8.739 | 0.0000 | [33;-88;-1] |
| | left postcentral | 0.0023 | 181 | 6.971 | 0.0000 | [-54;-25;44] |
| | left postcentral | | | 5.316 | 0.0000 | [-39;-31;48] |
| Semantic features | right temporal occipital fusiform | 0.0000 | 4151 | 11.149 | 0.0000 | [30;-49;-13] |
| | right inferior lateral occipital | | | 8.674 | 0.0000 | [33;-88;-1] |
| | right temporal occipital fusiform | | | 8.397 | 0.0000 | [24;-40;-20] |
| | left supramarginal | 0.0011 | 206 | 7.146 | 0.0000 | [-51;-34;36] |
| | left postcentral | | | 4.173 | 0.0004 | [-66;-19;29] |
| Semantic features (partial) | right temporal occipital fusiform | 0.0000 | 1268 | 12.072 | 0.0000 | [30;-49;-13] |
| | right temporal occipital fusiform | | | 8.203 | 0.0000 | [24;-40;-20] |
| | right temporal posterior fusiform | | | 7.363 | 0.0000 | [36;-34;-16] |
| | left temporal occipital fusiform | 0.0000 | 564 | 7.570 | 0.0000 | [-30;-55;-9] |
| | left lingual | | | 6.492 | 0.0000 | [-21;-58;-5] |
| | left inferior lateral occipital | | | 5.976 | 0.0000 | [-45;-67;-1] |
| | right precuneus | 0.0012 | 198 | 7.305 | 0.0000 | [6;-55;36] |
| | left precuneous | | | 4.747 | 0.0001 | [-9;-49;44] |
| | right precuneus | | | 4.329 | 0.0003 | [9;-64;25] |
| | left supramarginal | 0.0199 | 108 | 6.121 | 0.0000 | [-51;-34;36] |

Coordinates in MNI space, and region labels from the Harvard-Oxford Atlas

#### Table S2. RSA sensor effects for the partial correlation analysis

| *Model RDM* | *Model Peak (95% CIs)* | *Time window* | *cluster mass* | *cluster p* |
|---|---|---|---|---|
| CORnet-V1 | 100 (94-102) | '85-114 ms' | 54.53 | 0.0463 |
| | | '355-414 ms' | 89.13 | 0.0162 |
| CORnet-V2 | 120 (120-128) | '109-154 ms' | 93.93 | 0.0108 |
| CORnet-V4 | 96 (88-102) | '67-140 ms' | 133.42 | 0.0032 |
| CORnet-IT | 258 (256-298) | '131-602 ms' | 891.53 | <0.0001 |
| | | '607-646 ms' | 53.83 | 0.0462 |
| Semantic features | 352 (274-382) | '259-400 ms' | 217.76 | 0.0011 |
| | | '491-538 ms' | 79.22 | 0.0228 |
| | | '595-802 ms' | 396.77 | 0.0002 |

#### Table S3. RSA sensor effects for the semantic feature model controlling for different ANNs

| *Control model RDMs* | *Time window* | *cluster mass* | *cluster p* |
|---|---|---|---|
| Alexnet | '103-802 ms' | 1540.58 | 0.0001 |
| CORnet-S | '259-400 ms' | 217.76 | 0.0011 |
| | '491-538 ms' | 79.22 | 0.0228 |
| | '595-802 ms' | 396.77 | 0.0002 |
| CORnet-RT | '249-548 ms' | 627.15 | 0.0002 |
| | '581-802 ms' | 458.87 | 0.0003 |
| Resnet50 | '111-802 ms' | 1549.88 | <0.0001 |
| VGG19 | '103-174 ms' | 122.87 | 0.0103 |
| | '249-802 ms' | 1205.94 | <0.0001 |
| No control | '73-802 ms' | 2092.03 | <0.0001 |

#### Table S4. RCA results showing feedforward (FF) and feedback (FB) effects

| *Model RDM* | *Direction* | *Model Peak (95% CIs)* | *Time window* | *cluster mass* | *cluster p* |
|---|---|---|---|---|---|
| CORnet-V1 | FF | 110 (106-112) | '89-140 ms' | 114.86 | 0.0032 |
| | FF | | '177-218 ms' | 71.98 | 0.0156 |
| | FF | | '381-442 ms' | 97.84 | 0.0065 |
| | FF | | '577-612 ms' | 63.18 | 0.0209 |
| | FF | | '627-670 ms' | 70.35 | 0.0165 |
| CORnet-V2 | FF | 138 (132-138) | '117-166 ms' | 127.78 | 0.0003 |
| CORnet-V4 | FF | 112 (106-116) | '75-178 ms' | 287.30 | 0.0002 |
| | FF | | '237-388 ms' | 257.31 | 0.0003 |
| | FF | | '421-472 ms' | 76.01 | 0.0141 |
| | FF | | '537-582 ms' | 67.91 | 0.0195 |
| | FF | | '605-648 ms' | 71.10 | 0.0180 |
| CORnet-IT | FF | 260 (258-274) | '75-716 ms' | 1660.74 | <0.0001 |
| Semantic features | FF | 298 (298-318) | '273-802 ms' | 937.33 | <0.0001 |
| CORnet-IT | FB | 334 (262-374) | '243-408 ms' | 306.26 | 0.0003 |
| | FB | | '501-536 ms' | 53.75 | 0.0273 |
| Semantic features | FB | 398 (384-422) | '383-416 ms' | 52.71 | 0.0276 |
| | FB | | '523-552 ms' | 52.17 | 0.0285 |
| | FB | | '601-802 ms' | 358.31 | <0.0001 |

#### Table S5. RCA results for the semantic model RDM

| | Model peak (95% CI) | Time window | cluster mass | cluster p |
|---|---|---|---|---|
| pVTC to right ATL | 324 (300-326) | '147-802 ms' | 1338.48 | < 0.001 |
| pVTC to left PFC/ATL | 302 (290-310) | '147-416 ms' | 497.84 | < 0.001 |
| pVTC to left PFC/ATL | | '555-632 ms' | 109.14 | 0.009 |
| pVTC to left PFC/ATL | | '643-802 ms' | 235.69 | 0.001 |
| Right ATL to pVTC | 402 (386-448) | '385-462 ms' | 116.44 | 0.0058 |
| Left PFC/ATL to pVTC | 290 (284-470) | '441-496 ms' | 97.11 | 0.0071 |
| Left PFC/ATL to pVTC | | '601-656 ms' | 91.09 | 0.0090 |
| Left PFC/ATL to right ATL | 462 (390-472) | '439-482 ms' | 83.85 | 0.0078 |

#### Table S6. RCA results for visual and semantic model RDMs

| Model RDM | Direction | Peak time (95% CI) | Time window | cluster mass | cluster p |
|---|---|---|---|---|---|
| CORnet-V1 | pVTC to right ATL | 116 (114-116) | '93-154 ms' | 130.79 | 0.006 |
| | pVTC to right ATL | | '335-438 ms' | 160.85 | 0.003 |
| CORnet-V2 | pVTC to right ATL | 134 (134-134) | '103-216 ms' | 223.45 | 0.002 |
| CORnet-V4 | pVTC to right ATL | 114 (102-148) | '69-400 ms' | 772.08 | 0.000 |
| CORnet-IT | pVTC to right ATL | 278 (122-288) | '81-756 ms' | 1690.44 | 0.000 |
| Semantic features | pVTC to right ATL | 324 (300-326) | '147-802 ms' | 1338.48 | 0.000 |
| CORnet-V1 | pVTC to left PFC/ATL | 116 (110-118) | '93-150 ms' | 111.39 | 0.004 |
| CORnet-V2 | pVTC to left PFC/ATL | 136 (126-136) | '75-200 ms' | 192.36 | 0.001 |
| CORnet-V4 | pVTC to left PFC/ATL | 136 (104-136) | '75-268 ms' | 364.23 | 0.000 |
| | pVTC to left PFC/ATL | | '289-478 ms' | 338.55 | 0.000 |
| CORnet-IT | pVTC to left PFC/ATL | 118 (116-136) | '79-754 ms' | 1248.74 | 0.000 |
| Semantic features | pVTC to left PFC/ATL | 302 (290-310) | '147-416 ms' | 497.84 | 0.000 |
| | pVTC to left PFC/ATL | | '555-632 ms' | 109.14 | 0.009 |
| | pVTC to left PFC/ATL | | '643-802 ms' | 235.69 | 0.001 |
| CORnet-IT | Right ATL to pVTC | 282 (246-290) | '169-324 ms' | 312.79 | 0.0006 |
| | Right ATL to pVTC | | '347-418 ms' | 103.05 | 0.0103 |
| Semantic features | Right ATL to pVTC | 402 (386-448) | '385-462 ms' | 116.44 | 0.0058 |
| | Left PFC/ATL to pVTC | | '441-496 ms' | 97.11 | 0.0071 |
| | Left PFC/ATL to pVTC | | '601-656 ms' | 91.09 | 0.0090 |
| | Left PFC/ATL to right ATL | | '439-482 ms' | 83.85 | 0.0078 |
